# Identification of differentially expressed pyroptosis-related genes in LUAD

**DOI:** 10.1101/2022.08.27.505531

**Authors:** Bao Qian, Jiuzhou Jiang

## Abstract

**Background:** Lung cancer has the highest mortality rate among malignant tumors in the world, and adenocarcinoma, the most important pathological type of lung cancer, accounts for 80%-85% of all lung cancers, causing a significant burden of disease. Pyroptosis is a newly recognized form of programmed cell death (PCD).

**Methods:** 288 normal tissue data from The Genotype-Tissue Expression (GTEx) database and 501 tumor tissue data from The Cancer Genome Atlas (TCGA) database were selected. Differentially expressed genes were identified between normal and LUAD patients.

**Results and Conclusions:** With a consensus cluster analysis, 501 LUAD patients were divided into two clusters, and there was a significant difference in survival time between the two clusters. Ten differentially expressed genes were identified, with four up-regulated and six down-regulated. Pyroptosis is closely related to LUAD.

## 1. Introduction

With an estimated 2 million new cases and 1.76 million deaths per year, lung cancer is the second most prevalent cancer in the world [1, 2]. The most common histologic subtype of lung cancer, lung adenocarcinoma (LUAD), accounts for the majority of cancer deaths[2, 3]. Over the past decade, the treatment of NSCLC dramatically changed, mainly due to advances in biomarkers that have led to great success in patient-specific targeted therapies and immunotherapy [4]. Although rapid advances in diagnosis and treatments, 5-year overall survival in NSCLC remains poor, only 15% of LUAD patients survive beyond five years [5–7]. Therefore, it is essential to identify new prognostic genetic signatures.

Pyroptosis is a newly recognized form of programmed cell death (PCD) that is pro-inflammatory and distinct from iron-dead cell death, apoptosis, and autophagy[8]. The critical role of caspase 1 and caspase 11 in the lytic form of cell death is widely associated with protecting organisms from pathogens, called pyroptosis[9]. Pyroptosis is characterized by rapid plasma membrane rupture and the release of pro-inflammatory intracellular contents, in contrast to apoptosis, which is characterized by cellular content packaging and non-inflammatory phagocytic uptake by membrane-bound apoptotic vesicles[10, 11]. NLRP3[12], Gasdermin D (GSDMD)[13], Caspase 1 (CASP1)[14], etc., which are all Pyroptosis-related genes (PRG), are closely associated with tumorigenesis and tumor progression.

Next-generation sequencing (NGS) is a high-throughput gene sequencing tool that can identify various types of tumor-related genetic variants and is widely used for diagnosis and molecular typing, guiding therapeutic drug selection, genetic risk-related gene assessment, etc. [15] These genetic data can help patients obtain important tools for clinical tumor precision diagnosis and treatment.

Given the available findings, we know that there is a significant role for pyroptosis in tumor development and in the treatment of tumors. Therefore, studying genes related to pyroptosis is of great value for the assessment and prognosis of LUAD and may be an essential therapeutic trend for LUAD.

## Method

### Data acquisition and processing

Gene expression data (count matrix), corresponding clinical information, and the somatic mutation profiles for 585 LUAD patients were downloaded from The Cancer Genome Atlas (TCGA) via the UCSC Xena browser (https://xenabrowser.net/). In this study, we kept the expression profiles of the primary solid tumor samples, removed patients with missing survival information, and filtered out patients with less than 30 days of follow-up. 501 LUAD patients were included in the analysis. Furthermore, we downloaded RNA-seq data from the GTEx database (https://xenabrowser.net/datapages/) for 288 normal human lung samples. For RNA-seq data, the TPM algorithm was used to convert the FKPM data into transcript per million (TPM), which was used to estimate each gene’s expression level before log2 transformations were performed. Based on the mutation data, the “Maftools” package (V 3.38.1) was used to analyze these results. Moreover, the differential expression of genes in the normal and cancer groups was shown by a box plot.

### Analysis of pyroptosis-related genes

We conducted a consensus clustering analysis using the *ConsensusClusterPlus* (V1.60.0) on all 501 LUAD patients in the TCGA cohort to explore the association between the expression of 31 pyroptosis-associated genes and LUAD subtypes. The parameters were as follows: reps = 100, pitem = 1, pfeature = 0.8, distance = “Spearman”, clusterAlg = “hc”, innerLinkage = “ward.D2”, finalLinkage = “ward.D2”.

### Identification of differential expression genes

Based on the previous studies, 33 pyroptosis-related genes were selected[16]. Normal and tumor tissues were selected for differential expression analysis using limma with threshold: FDR < 0.05 and log2| FC | > 1.5. GO and Kyoto Encyclopedia of Genes and Genomes (KEGG) enrichment analysis was then performed on selected differentially expressed genes (DEGs), showing the results with FDR < 0.05.

## Results

### Analysis of pyroptosis-related genes in LUAD

Genetically, mutations in cell pyroptosis-related genes were present in 209 of 616 samples (approximately 33.93%, Figure 1A). We also investigated the association between pyroptosis-related genes with CNV and chromosomes (Figure 1B). To compare the expression levels of pyroptosis-related genes in 288 normal tissues from the GTEx database and 501 tumor tissue data from TCGA, we calculated the expression levels of 31 genes related to pyroptosis. Figure 1C demonstrates the differential expression of 31 pyroptosis-related genes in normal versus LUAD patients.

**Figure 1.**
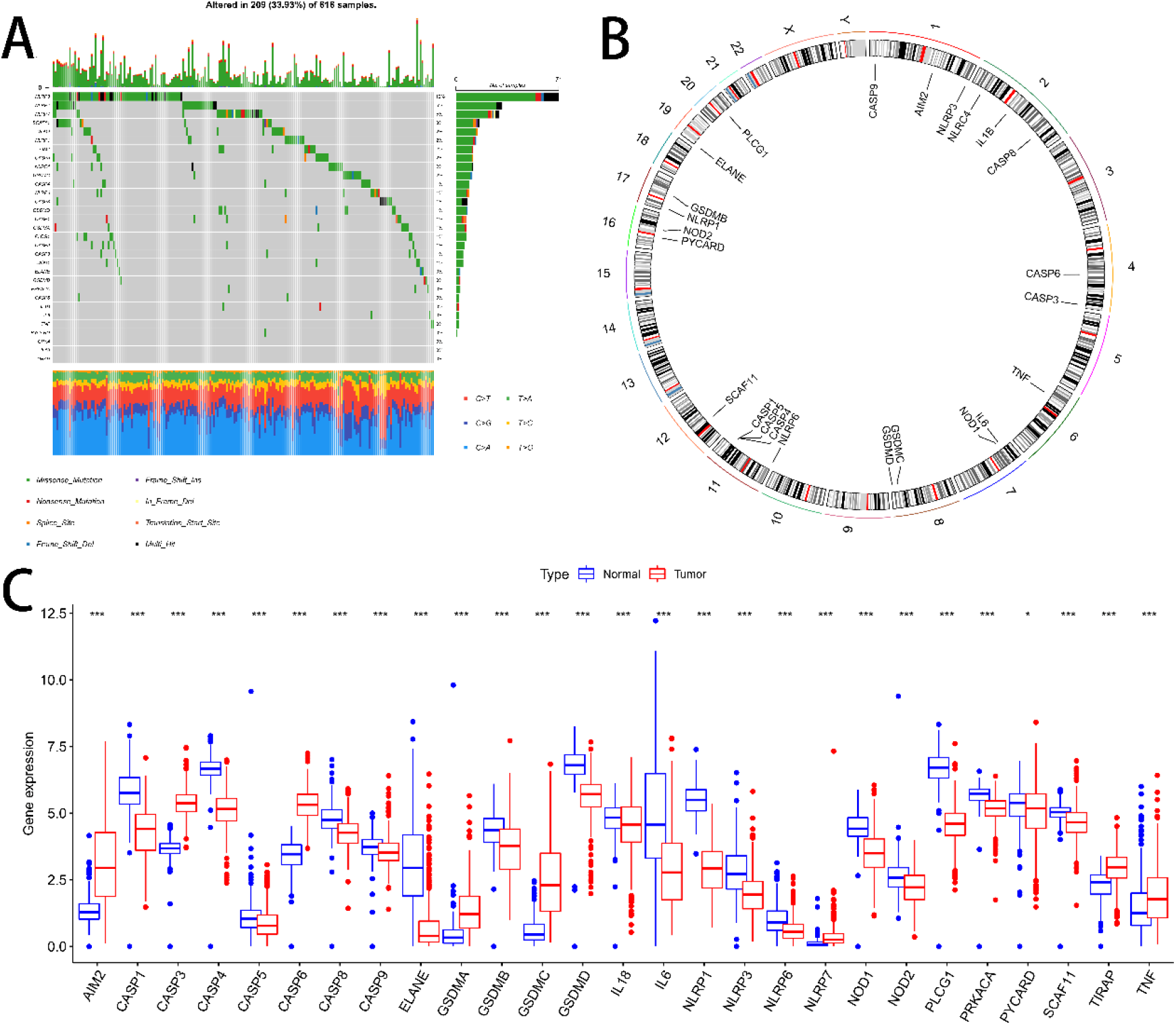
(A) The association between pyroptosis-related genes with CNV and chromosomes. (B) Waterfall plot displaying details of mutations in pyroptosis-related genes. (C) Box plot of pyroptosis-related genes, red represents the tumor group, blue represents the normal group.

### Tumor classification based on the pyroptosis-related genes

A consensus clustering analysis was performed on all 501 LUAD patients in the TCGA cohort for the purpose of exploring the association between the expression of 31 pyroptosis-related genes and LUAD subtypes. We found the highest intra-group and low inter-group correlations at k = 2, suggesting that LUAD patients may be well divided into two clusters (Figure 2A). We also examined the OS time of C1 versus C2 and found a significant difference (p = 0. 034, Figure 2B). Figure 2C shows the expression of pyroptosis-related genes in the two clusters.

**Figure 2.**
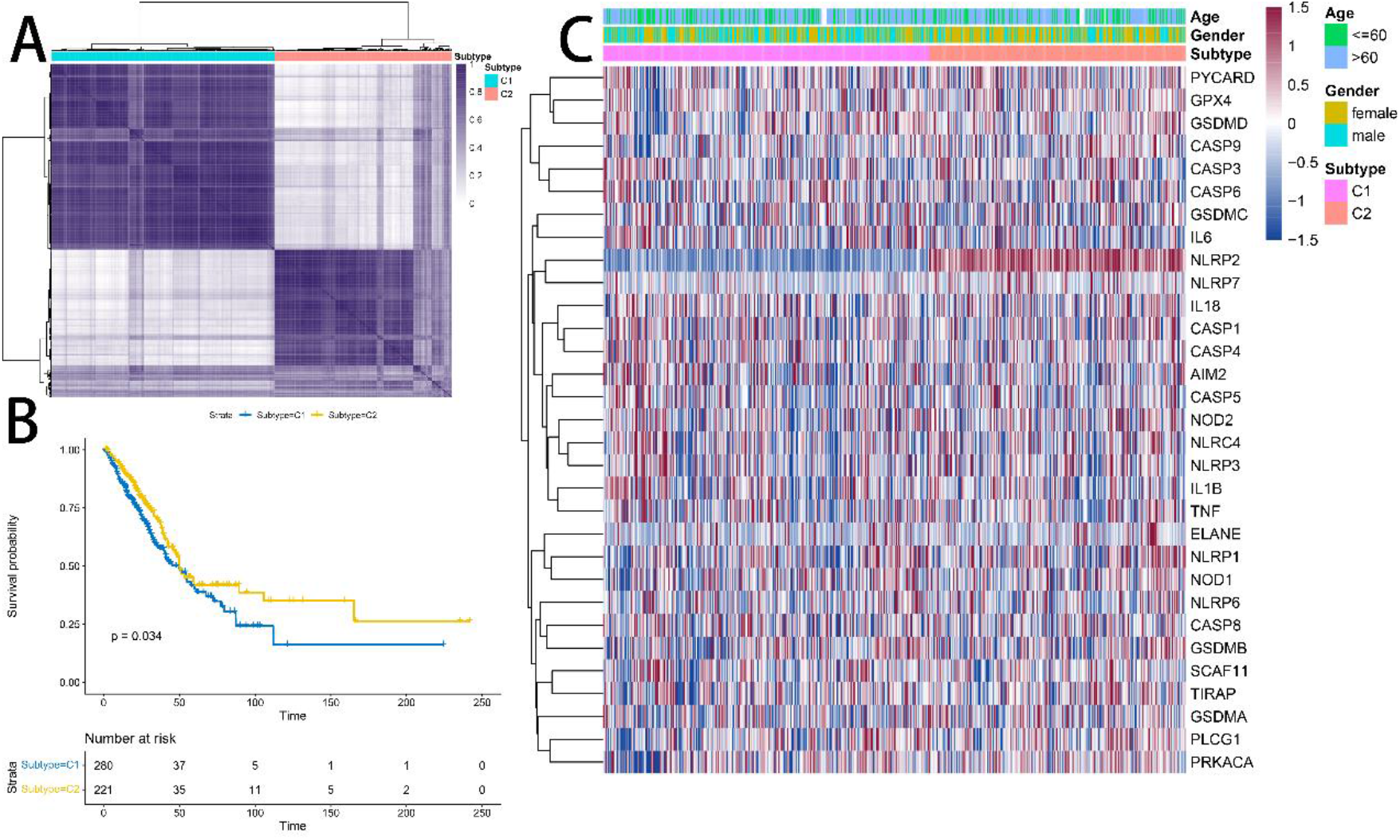
(A) Consensus clustering matrix. 501 LUAD patients were divided into two clusters. (B) Kaplan-Meier survival curves for C1 and C2. The clinical outcome endpoint was OS. (C) Heatmap of clinical features and gene expression changes between normal and cancer groups.

### Differential expression analysis between normal and LUAD groups

For further examination of the impact of these genes on cancer prognosis, the “limma” package (V3.52.2) was used to identify 10 DEGs, of which four genes (AIM2, GSDMC, CASP3, CASP6) were up-regulated and six (IL6, ELANE, CASP4, CASP1, NLRP1, PLCG1) genes were down-regulated (Figure 3A). Figure 3B shows a heat map of the up- and down-regulated genes. In order to further explore the function of pyroptosis-related genes, we performed GO enrichment analysis and KEGG pathway analysis on 10 DEGs, and the annotation results (FDR<0.05) are visualized in Figure 3C-3D. The results showed that pyroptosis, regulation of inflammatory response, inflammasome complex, and endopeptidase activity in cancer and cell cycle were significantly enriched in GO analysis. KEGG pathway analysis indicated that genes related to pyroptosis are associated with apoptosis, NOD−like receptor signaling pathway, and neutrophil extracellular trap formation. Figure 3E illustrates the single-gene Kaplan Meier curve. The findings indicated a significant difference in GSDMC, CASP3, CASP6, ELANE, CASP4, CASP1, NLRP1, and PLCG1 (P < 0.05). There was no significant difference between AIM2 and IL6 (P > 0.05).

**Figure 3.**
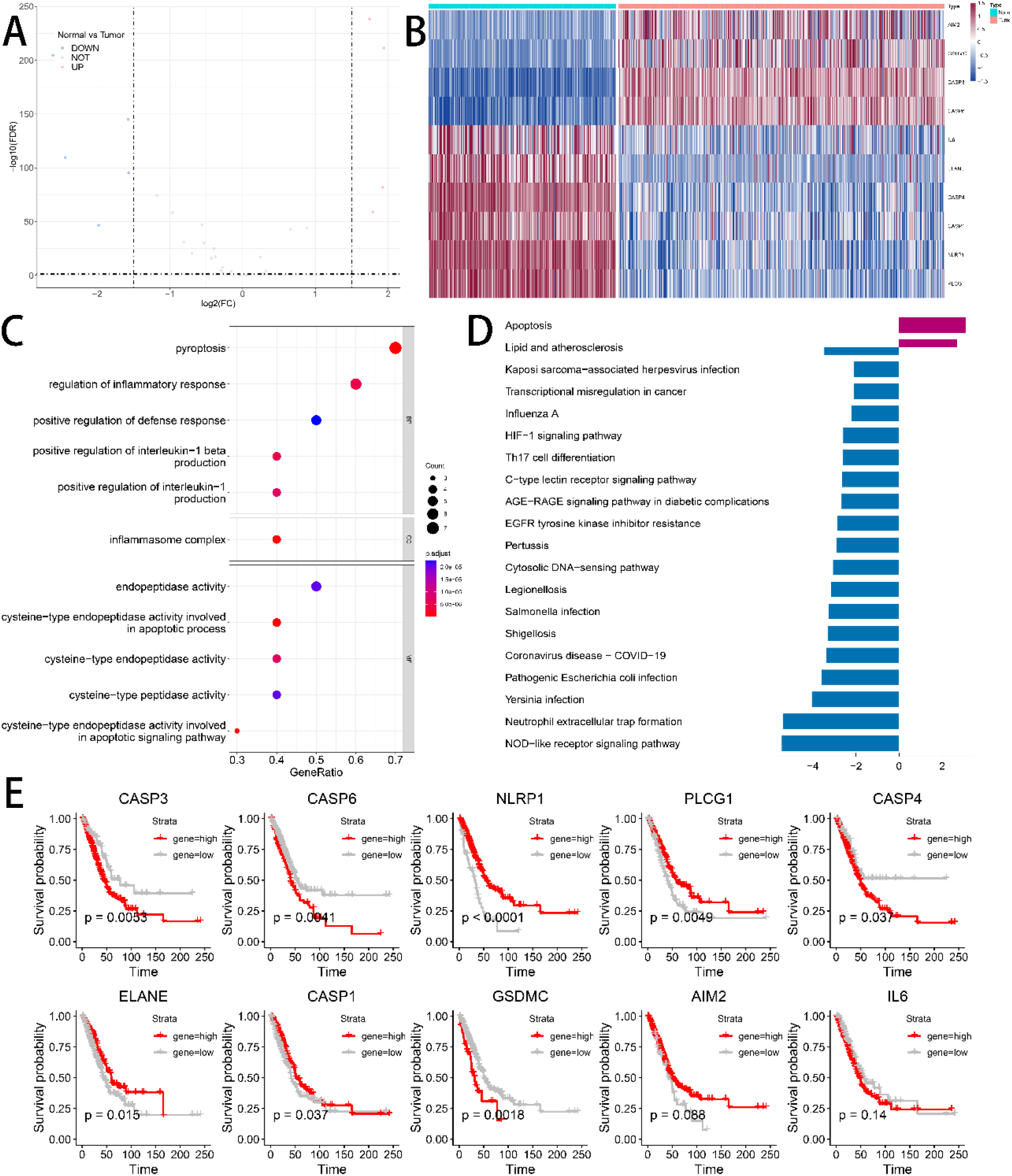
(A) Volcano map of differentially expressed genes for normal and LUAD patients. (B) Heatmap of differentially expressed genes in the TCGA dataset for normal and LUAD patients. (C) Bubble plot of annotation using GO for differentially expressed genes. (D) Annotation using KEGG for differentially expressed genes. (E) Single gene Kaplan Meier curve.

## Discussion

Lung cancer has the highest mortality rate among malignant tumors in the world, and adenocarcinoma, the most important pathological type of lung cancer, accounts for 80%-85% of all lung cancers, causing a significant burden of disease[17]. China has a high incidence of lung cancer, with 787,000 new cases of lung cancer and 631,000 deaths due to lung cancer in 2015 alone, accounting for the highest number of new incidences and deaths of malignant tumors in 2015[18]. From the information in the SEER database, the 5-year survival rate for lung cancer is about 26%.

Therefore, early prognostic assessment of lung adenocarcinoma and individualized treatment to reduce metastasis, recurrence, and prolong survival is particularly important. Therefore, the search for new prognostic markers becomes particularly important.

The primary function of pyroptosis, a process of inflammatory cell death, is to induce a robust inflammatory response and protect the host from microbial infection[19]. Inflammatory vesicles induce pyroptosis in two ways, referred to as canonical and noncanonical inflammatory vesicle pathways. As a new form of programmed cell death, pyroptosis has been studied by many scholars in recent years and has been proposed as a target for lung cancer treatment by affecting pyroptosis[13, 20].

In recent years, as cancer research has become more and more advanced, the research on new biomarkers and prognostic markers of lung cancer is also increasing[21–23]. These provide direction for the prognosis of lung cancer patients. Several studies have associated the role of pyroptosis-related genes in the pathogenesis and prognosis of lung cancer, highlighting the clinical importance of pyroptosis in the therapeutic approach to lung cancer[24].

In the present study, we collected 33 pyroptosis-related genes using the results of previous studies and performed several analyses. In the TCGA and GTEx database, 31 out of 33 genes were matched, and 27 genes were significantly differentially expressed between normal and cancer groups. The *ConsensusClusterPlus* package was used to conduct consensus clustering analysis on 501 LUAD patients. Patients were well divided into two groups, with significant differences in survival analysis between the groups. We analyzed ten differentially expressed genes, with four up-regulated and six down-regulated.

There are also some limitations to this study. Firstly, the regulatory mechanism and network of pyroptosis are not completely clear and need to be further explored. We may not have included the full range of pyroptosis-related genes for analysis because of the limited understanding of pyroptosis.

## Conclusions

With a consensus cluster analysis, 501 LUAD patients were divided into two clusters, and there was a significant difference in survival time between the two clusters. Ten differentially expressed genes were identified, with four up-regulated and six down-regulated. Pyroptosis is closely related to LUAD.

## Notes

### Competing Interest Statement

The authors have declared no competing interest.

